# Contributions from Long-Term Memory Explain Superior Visual Working Memory Performance with Meaningful Objects

**DOI:** 10.1101/2025.07.07.663569

**Authors:** Hyung-Bum Park, Edward Awh

**Affiliations:** Institute for Mind and Biology, University of Chicago Department of Psychology, University of Chicago 940 East 57th St., Chicago, IL 60637

## Abstract

Visual working memory (WM) capacity has been claimed to be larger for meaningful objects than for simple features, possibly because richer semantic representations enhance the distinctiveness of stored items. However, prior demonstrations typically compared trial-unique meaningful objects with a small set of repeated simple features. This design confounds meaningfulness with proactive interference (PI), such that PI is minimal for trial-unique objects but substantial for repeated features. As a result, superior performance for meaningful objects may reflect contributions from episodic long-term memory (LTM) rather than expanded WM capacity. To test this, Experiment 1 measured WM for repeated colors, repeated meaningful objects, and trial-unique meaningful objects. The advantage for objects over colors was replicated in the trial-unique condition, but eliminated for repeated objects that equated PI across stimulus types. Hierarchical Bayesian dual-process modeling revealed that the trial-unique advantage reflected stronger familiarity signals, whereas recollection remained stable across stimulus types. Experiment 2 assessed WM storage directly using contralateral delay activity (CDA), an electrophysiological marker of the number of items stored. Although trial-unique objects again yielded behavioral advantages, CDA activity across increasing set sizes revealed a common slope and plateau for trial-unique meaningful objects and repeated colors. The CDA difference between stimulus types was additive and did not vary with set size, providing no evidence for an increased number of stored items. These findings demonstrate that previously reported advantages for meaningful objects primarily reflect reduced PI and enhanced LTM familiarity. When PI is equated, WM storage limits for simple and meaningful stimuli are equivalent.

**Significance Statement:** Working memory provides the mental workspace that underlies reasoning, learning, and everyday decision making, yet its capacity is sharply limited. Previous studies suggested that meaningful, real-world objects are remembered better, raising the possibility that knowledge can expand this known capacity limit. However, many designs confound working memory with long-term familiarity. Here, equating proactive interference removed the behavioral advantage for meaningful items. A neural marker of active storage showed additive differences between stimulus types that did not vary with load, indicating no increase in the number of stored items. These findings identify interference, rather than expanded storage, as the source of the reported advantage and offer practical guidance for future experimental design and theories of memory limits.

---

Visual working memory (WM) temporarily maintains visual information to support goal-directed behavior (Cowan, 2001; Luck & Vogel, 2013). This online system is highly limited in capacity, such that performance drops sharply when memory arrays contain more than three-or-four items (Alvarez & Cavanagh, 2004; Awh et al., 2007; Luck & Vogel, 1997; Ngiam et al., 2023; Park et al., 2017; Zhang & Luck, 2008; Zhao et al., 2023). While some models argue against strict limits (Bays, 2015; Schurgin et al., 2020; van den Berg et al., 2012, 2014), most work converges on the conclusion that WM supports only a few individuated items with *sufficient fidelity* to guide behavior (Adam et al., 2017; Oberauer & Lin, 2017; Park & Zhang, 2024).

In contrast, recent reports of superior WM performance for meaningful, real-world objects challenge this item-based fixed-capacity view (Asp et al., 2021; Brady et al., 2016; Chung et al., 2024; Torres et al., 2024). Such advantages have been attributed to semantic knowledge in long-term memory (LTM) that promotes deeper encoding and distinctive representations, potentially allowing more items to be maintained concurrently (Brady & Alvarez, 2011; Curby et al., 2009). However, most studies employed trial-unique objects, each presented only once, and compared performance to a repeated set of simple features (e.g., categorically-distinct colors^1^). This design confounds meaningfulness with proactive interference (PI; Jonides & Nee, 2006; Kane & Engle, 2000). Because PI is virtually absent with trial-unique stimuli, recognition can be guided by global familiarity, which supports surprisingly large capacity estimates (Endress & Potter, 2014; Standing, 1973). Thus, meaningful-object advantages may be driven by reduced PI affording stronger contributions from LTM rather than expanded WM storage.

Neural measures provide a complementary test. The contralateral delay activity (CDA) is a well-established marker tracking the number of items stored in WM (Luria et al., 2016; Vogel & Machizawa, 2004). CDA amplitude scales with the *number* of items maintained, not the amount of information per item. Recent studies have reported larger CDA amplitudes for meaningful compared to simple stimuli and interpreted this as evidence for greater WM storage (Brady et al., 2016; Thibeault et al., 2024; Torres et al., 2024). Because they typically measured CDA at only a single set size^2^, however, these designs cannot rule out additional effects unrelated to load. For example, Woodman and Vogel (2008) found equivalent CDA amplitudes for four orientations versus four color-orientation conjunctions, despite the latter containing twice as many features. Importantly, they also showed orientations and conjunctions produced larger CDA amplitudes than colors, but these differences were *additive* with set size (Figure 1). If increased CDA amplitude reflected increased storage-related activity, this effect should increase as more items are stored. Thus, these additive stimulus-driven effects do not appear to reflect WM storage, per se. Similarly, Drew and colleagues (2011) found increased CDA amplitude during multiple-object tracking relative to static WM tasks, but this effect was also additive with set size, suggesting a stimulus-driven effect unrelated to WM storage. These findings make clear that CDA must be examined across multiple set sizes to discriminate between changes in WM storage and stimulus-driven effects that do not determine the number of items stored (Quirk et al., 2020).

**Figure 1.**
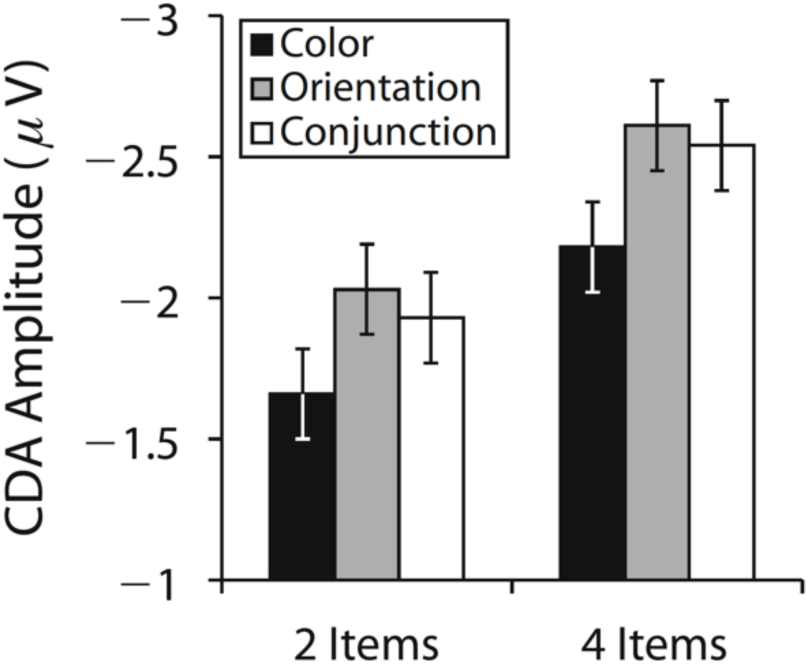
Mean contralateral delay activity (CDA) amplitudes for color, orientation, and conjunction stimuli at set sizes 2 and 4, adapted from. Figure 3D **of Woodman and Vogel (2008).** Although orientation and conjunction stimuli elicited larger CDA amplitudes than color stimuli, these differences were additive across set sizes, suggesting that the enhanced CDA was not due to an increased number of items stored in working memory.

The present study addresses this issue with converging behavioral and neural measures. Experiment 1 equated PI by comparing recognition performance across repeated colors, repeated meaningful objects, and trial-unique meaningful objects. Confidence-based judgments were modeled with a hierarchical Bayesian dual-process signal detection (DPSD) framework to separate recollection, reflecting context-bound WM, from episodic familiarity (Yonelinas, 2002; Yonelinas et al., 2010). Experiment 2 measured CDA across multiple set sizes for trial-repeated colors and trial-unique meaningful objects in a lateralized change-detection task. If meaningful objects expand WM capacity, advantages should persist under equated PI, and CDA functions should diverge across stimulus types, producing a load-by-stimulus interaction. Conversely, if advantage reflects episodic familiarity under reduced PI, CDA differences should be *additive* across set sizes, consistent with stimulus-driven rather than storage-related effects.

## Experiment 1

To test whether superior visual WM performance for meaningful objects reflects enhanced WM capacity or reduced PI, Experiment 1 independently manipulated meaningfulness and PI. We compared recognition performance across three stimulus conditions: repeated colors, repeated meaningful objects, and trial-unique meaningful objects. If the meaningful object benefit reflects a genuine expansion of WM capacity, it should persist regardless of stimulus repetition. Conversely, if the benefit is due to the absence of PI with trial-unique stimuli, the advantage should disappear in the repeated meaningful object condition that equates PI. To further evaluate contributions of WM and episodic LTM, we analyzed participants’ confidence judgments using hierarchical Bayesian DPSD modeling, allowing separate estimates of recollection (context-bound WM) and familiarity (episodic LTM) signals.

## Materials and Methods

### Participants

Thirty-one volunteers (18 female) participated in the experiments and received monetary compensation ($20 per hour). Participants were aged between 18 and 31 years (*M* = 22.3, *SD* = 3.2), reported normal or corrected-to-normal visual acuity, and provided informed consent according to procedures approved by the University of Chicago Institutional Review Board.

Sample size was determined via an a priori power analysis (G*Power 3.1; Faul et al., 2009), targeting 85% power to detect a medium effect size (Cohen’s *d* > 0.5) in primary comparisons between stimulus conditions at an alpha level of 0.05, based on prior studies investigating visual WM capacity differences between meaningful objects and color stimuli (Brady et al., 2016; Chung et al., 2023, 2024).

### Experimental Design

Stimuli were generated using MATLAB (The MathWorks, Natick, MA, USA) and the Psychophysics toolbox (Brainard, 1997), and presented on an LCD computer screen (BenQ XL2430T; 120 Hz refresh rate; 61 cm screen size in diameter; 1920 × 1080 pixels) with a grey background (15.1 cd/m^2^) and positioned approximately 70 cm from participants.

Figure 2 illustrates the stimuli and procedure of the task. Participants performed a visual WM recognition task with a set size of six items across three blocked conditions: trial-repeated colors, trial-repeated meaningful objects, and trial-unique meaningful objects. Block order was counterbalanced across participants. Each block contained 120 trials. Participants were given short breaks between blocks and 20-second breaks after every 40 trials within each block to reduce fatigue.

**Figure 2.**
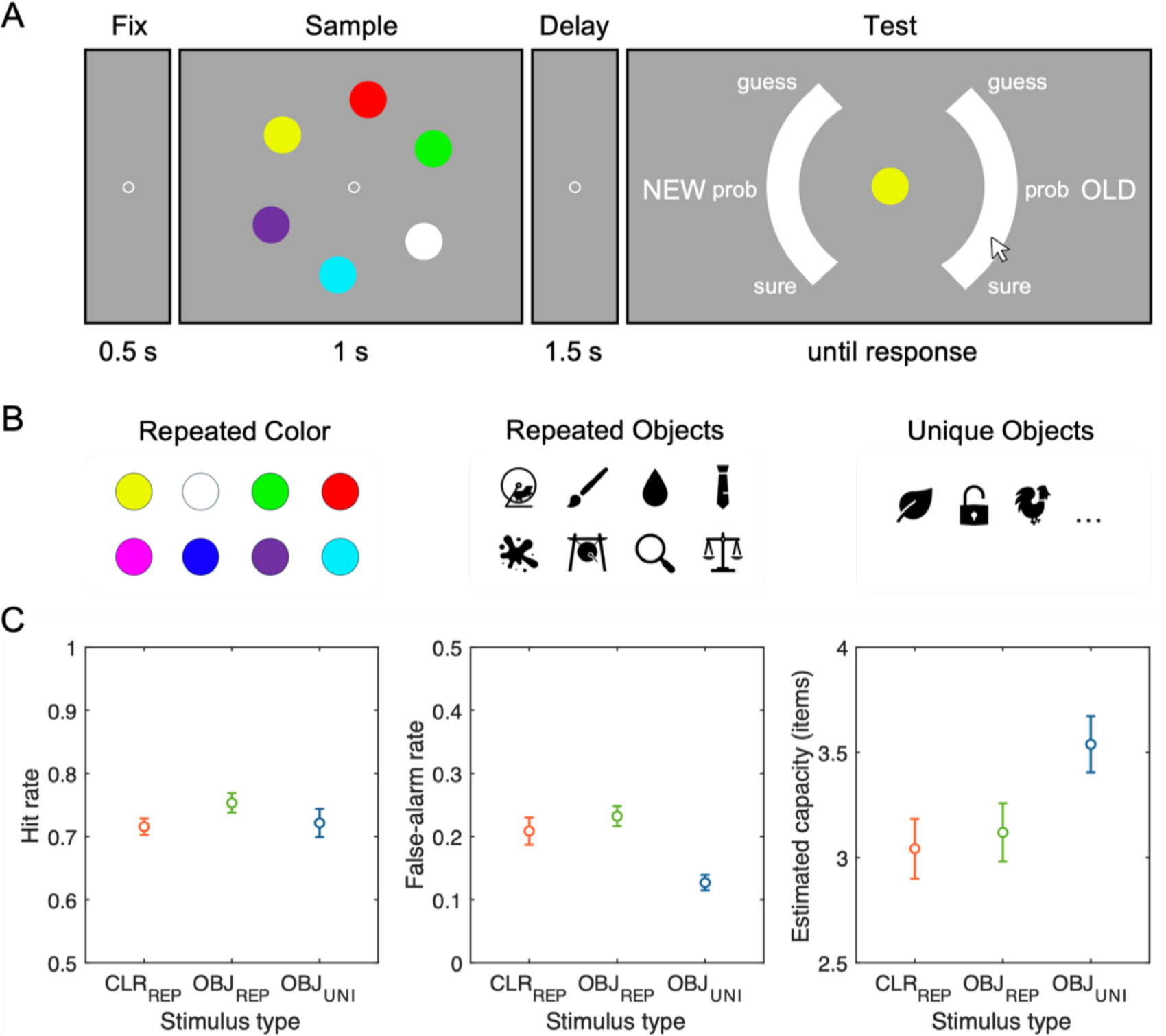
Experimental paradigm and behavioral results from Experiment 1. (A) Task sequence for the visual WM recognition task. Each trial began with a fixation display (500 ms), followed by a memory array containing six items (1,000 ms), a blank retention interval (1,500 ms), and a recognition probe requiring a confidence rating on a continuous scale ranging from “sure new” to “sure old”. (B) Examples of stimulus sets used across three conditions: colors (CLR_REP_; repeated across trials), meaningful objects repeated throughout the experiment (OBJ_REP_), and trial-unique meaningful objects presented only once (OBJ_UNI_). (C) Behavioral performance summarized as hit rate (left), false-alarm rate (middle), and estimated visual working memory capacity measured by Cowan’s *K* (right) for each stimulus condition. Error bars represent standard error of the means.

Each trial started with a 500 ms central fixation cross (0.2° visual angle), followed by a memory array of six items (each 2.0° × 2.0°) presented for 1,000 ms, arranged evenly along an imaginary circle (radius of 5.3°). After a 1,500 ms blank retention interval, participants saw a single central probe, with a semi-circular confidence rating scale (radius of 8.2°, thickness of 2.2°) split into left and right sides, separated by 90° gaps and with response regions labeled “NEW” and “OLD”, respectively. Participants responded by moving a mouse cursor from the screen center to the continuous scale labeled with discrete confidence markers, “surely-new”, “probably-new”, “guess-new”, “guess-old”, “probably-old”, and “surely-old”. Probe items were old (presented in the memory array) or new with equal probability.

In both trial-repeated conditions (colors and meaningful objects), memory arrays consisted of six items randomly selected from a fixed set of eight stimuli unique to each condition. The color set consisted of eight categorically distinct hues (RGB values: red [255,0,0], green [0,255,0], blue [0,0,255], magenta [255,0,255], yellow [255,255,0], cyan [0,255,255], orange [255,128,0], and white [255,255,255]). For objects, we selected Microsoft Office icons that were easily recognizable, including animals, plants, foods, transportation, musical instruments, and so forth (see Figure 2B for example stimuli). These items were presented in black on the grey background. The eight objects in the trial-repeated object condition were randomly sampled from the object pool for each participant to minimize potential influence of sampling bias in perceptual or semantic properties. The new probes were selected from the remaining items in the set of eight that were not presented in the current memory array. In contrast, for trial-unique objects condition, six novel objects were randomly selected each trial, with another previously unseen object used as the new probe.

### Statistical Analysis

#### Behavioral Performance

We calculated hit rates (correctly recognition of old probes) and false alarm rates (incorrect recognition of new probes as old) for each condition. From these, we computed Cowan’s *K* (capacity) = set size × (hit rate - false alarm rate).

#### Hierarchical Bayesian Dual-Process Signal Detection Modeling

Confidence ratings were derived from the final clicking response along the semi-circular response regions and binned into six discrete confidence levels, specifically, three levels of confidence for “old” responses (surely-old, probably-old, and guess-old) and three levels for “new” responses (guess-new, probably-new, and surely-new). These ratings were used to construct receiver operating characteristic (ROC) curves, plotting cumulative false-alarm rates on the x-axis against cumulative and hit rates on the y-axis across confidence levels.

These empirical ROCs were fitted with the DPSD model. The DPSD model assumes that recognition decisions are based on two independent processes (Yonelinas et al., 2010; Yonelinas, 2002): 1) Recollection, which is an all-or-none retrieval of contextual details reflecting context-bound WM representation, and 2) Familiarity, which is a continuous signal-detection process reflecting context-free episodic LTM traces. The DPSD parameter estimation was performed using a hierarchical Bayesian method, which provides robust population-level estimates of the model parameters by simultaneously accounting for different sources of uncertainty across individual (random) and condition (fixed) effects. The advantage of the hierarchical Bayesian method is especially useful in ROC modeling, wherein a relatively limited number of trials are allocated for each decision criterion (e.g., six-binned confidence level) per experimental condition (Park et al., 2023). The main and interaction effects were estimated in a general linear model, sampled from the normal distribution where the mean is the sum of the fixed and random effects and the variability term is the interaction across effects.

We used Markov chain Monte Carlo sampling to estimate the posterior distributions of the parameters, generating 12,000 samples after 12,000 warm-up iterations. Population-level parameter posteriors were thus represented by a 12,000 × 31 × 3 matrix (i.e., samples × participants × stimulus types). We chose noninformative and reasonably informative priors.

Model convergence was confirmed by using the Gelman–Rubin diagnostic *R^* (Gelman & Rubin, 1992), and found to be close to or equal to 1.0 for all population-level parameters. Figure 3 provides visualization of model fits over empirical ROCs for individual participant. Statistical inference was made based on posterior mean parameter estimates and their associated 95% highest density intervals (HDI95%). HDIs provide direct estimates of evidence strength, intervals that exclude zero provide strong evidence for credible effects (Kruschke, 2015).

**Figure 3.**
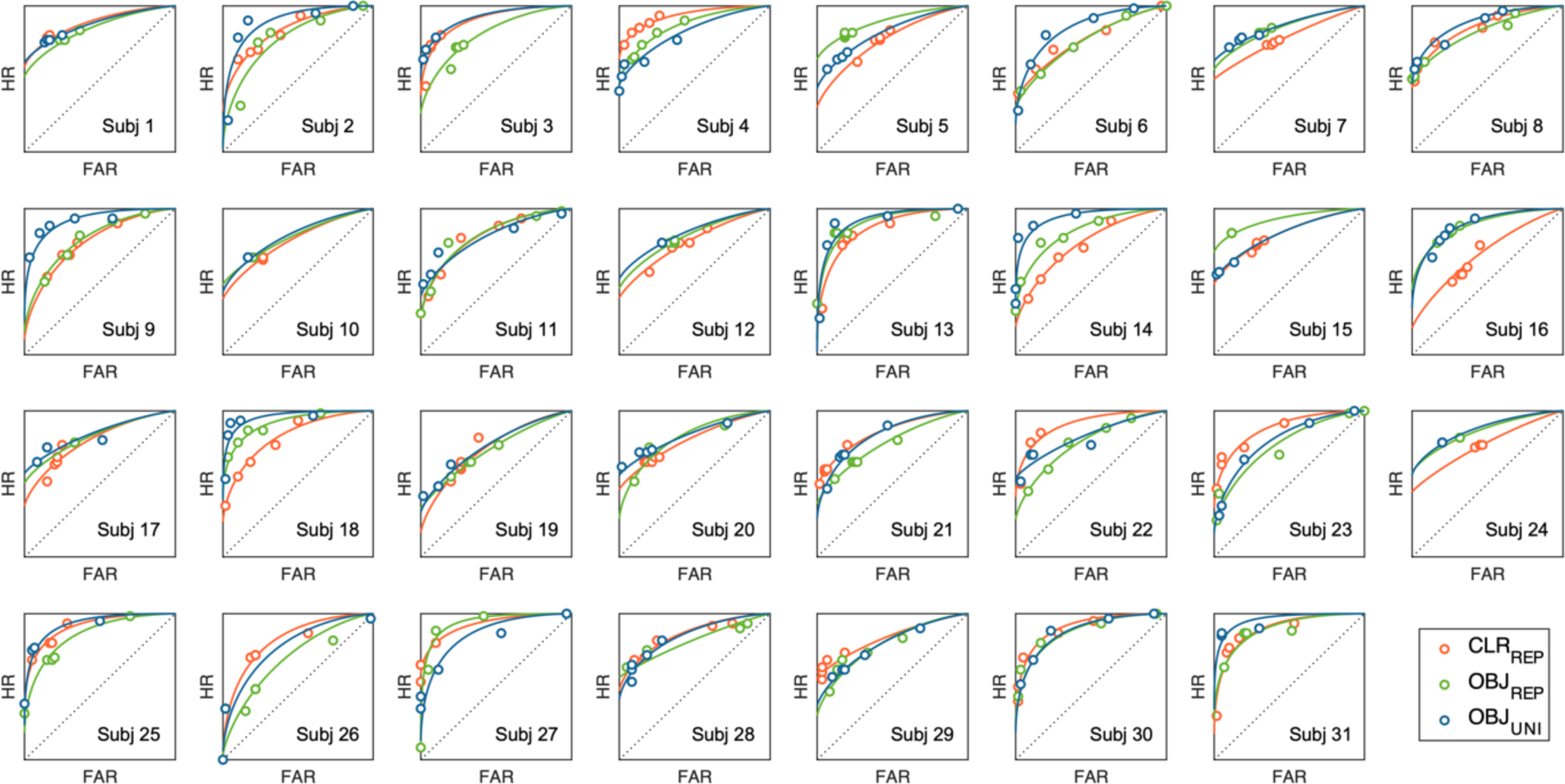
Observed and model-fitted ROC curves for individual participants. Each panel shows hit rate (HR) plotted over false-alarm rate (FAR) across confidence levels from each participant. Solid curves represent model predictions generated by the mean posterior parameters estimated from the hierarchical Bayesian dual-process signal detection model. Color codes represented repeated-colors (CLR_REP_ in red), repeated-objects (OBJ_REP_ in green), and unique-objects (OBJ_UNI_ in blue) conditions, respectively. Circles denote observed data. The close correspondence between predicted and observed values indicates good individual-level model fit.

## Code Availability

All data generated or analyzed during this study are available via the Open Science Framework repository at https://osf.io/e8ptm/

## Results

### Behavioral Performance

Figure 2C summarizes the hit rates, false alarm rates, and Cowan’s *K* values across the three experimental conditions. First, a one-way repeated-measures analysis of variance (ANOVA) on the mean Cowan’s *K*s revealed a significant main effect of stimulus type, *F*(2, 90) = 3.74, *p* = .028, *η^2^p* = .08. This effect was found to be primarily driven by higher capacity estimates for trial-unique objects (*M* = 3.54, *SD* = 0.75) compared to both trial-repeated colors (*M* = 3.04, *SD* = 0.79) and trial-repeated objects (*M* = 3.12, *SD* = 0.77), *ts*(30) > 2.69, Bonferroni-corrected *psbonf* < .036, *ds* > 0.49. Critically, Cowan’s *K* estimates did not significantly differ between trial-repeated colors and trial-repeated objects, *t*(30) = –0.42, *pbonf* = 1.000, *d* = –0.08.

Further analysis of hit and false alarm rates revealed that the observed advantage for trial-unique objects primarily stemmed from its reduced false alarms rather than enhanced hit rates. Specifically, for hit rates, a one-way repeated-measures ANOVA indicated no significant effect of stimulus type, *F*(2, 90) = 1.36, *p* = .263, *η^2^p* = .03, with highly comparable performance for trial-repeated colors (*M* = 0.72, *SD* = 0.07), trial-repeated objects (*M* = 0.75, *SD* = 0.09), and trial-unique objects (*M* = 0.72, *SD* = 0.13), *ts*(30) < 1.93, *psbonf* > .188, *ds* < 0.35. In contrast, false alarm rates varied significantly by stimulus type, *F*(2, 90) = 10.61, *p* < .001, *η^2^p* = .19, with trial-unique objects showing substantially fewer false alarms (*M* = 0.13, *SD* = 0.07) compared to both trial-repeated colors (*M* = 0.21, *SD* = 0.12) and trial-repeated objects (*M* = 0.22, *SD* = 0.09), *ts*(30) > 3.85, *psbonf* < .002, *ds* > 0.70. False alarm rates did not differ significantly between trial-repeated colors and trial-repeated objects, *t*(30) = 1.06, *pbonf* = .889, *d* = 0.19.

### Dual-Process Signal Detection Modeling

The hierarchical Bayesian DPSD model revealed a clear dissociation between recollection and familiarity processes across stimulus types (Figure 4). Recollection estimates were highly comparable across all three conditions, with substantial overlaps in their posterior distributions between trial-repeated colors (*M* = 0.35, *HDI95%* [0.33, 0.37]), trial-repeated objects (*M* = 0.35, *HDI95%* [0.32, 0.37]), and trial-unique objects (*M* = 0.36, *HDI95%* [0.34, 0.39]). In contrast, familiarity estimates exhibited a robust effect consistent with the results of Cowan’s *K*. Familiarity was credibly higher for trial-unique objects (*M* = 1.10, *HDI95%* [1.06, 1.15]) compared to both trial-repeated colors (*M* = 0.93, *HDI95%* [0.88, 0.98]) and trial-repeated objects (*M* = 0.91, *HDI95%* [0.85, 0.97]), with non-overlapping credible intervals providing strong evidence for this difference.

**Figure 4.**
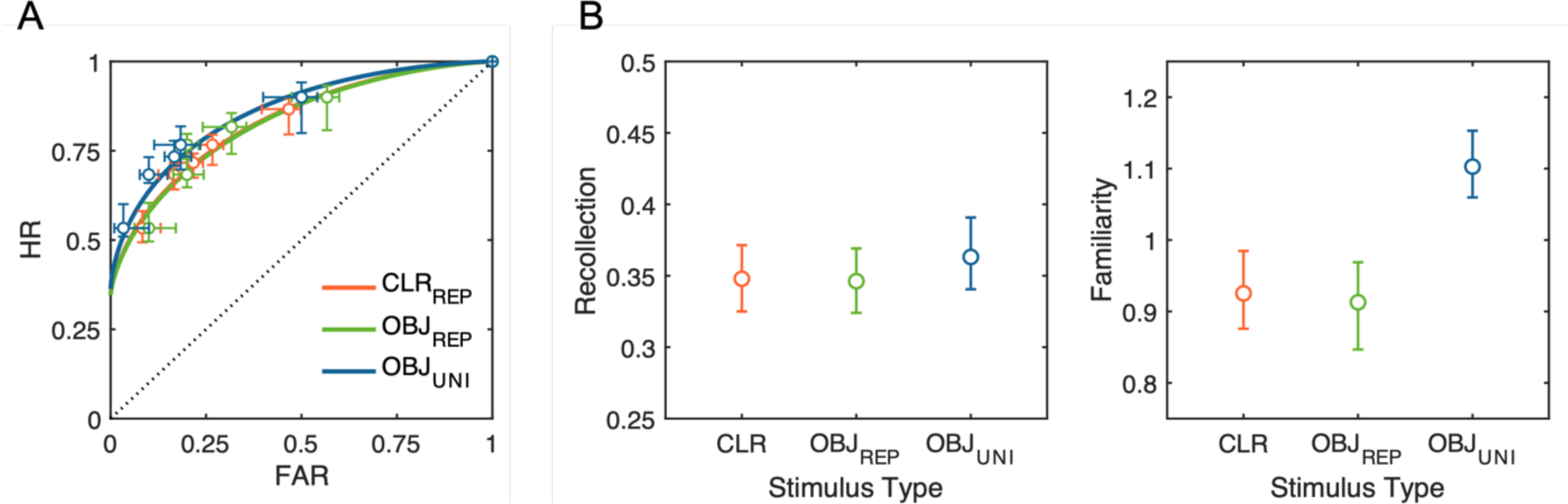
Observed and model-predicted ROC curves and posterior parameter estimates from the hierarchical Bayesian DPSD model. (A) ROC curves for each stimulus condition: repeated colors (CLR_REP_; red), repeated meaningful objects (OBJ_REP_; green), and trial-unique meaningful objects (OBJ_UNI_; blue). Circles represent observed mean hit rates (HR) and false-alarm rates (FAR) across confidence levels, and the horizontal and vertical error bars indicate standard errors of the mean FAR and HR, respectively. Solid lines depict model-predicted ROC curves based on posterior mean values of the model parameters. (B) Posterior means and 95% highest density intervals (HDIs_95%_) for the population-level recollection (left) and familiarity (right) parameters, across stimulus types. The boundaries of HDI_95%_ not crossing over between conditions indicate a statistically credible difference.

## Discussion

Experiment 1 demonstrated that improved memory performance for meaningful relative to simple objects may reflect reduced PI with trial-unique objects rather than increased WM capacity for meaningful items. Replicating past studies, we observed a clear performance advantage for meaningful objects, but only when the design employed trial-unique stimuli (Endress & Potter, 2014). This meaningful object benefit was eliminated when PI susceptibility was equated through repeated stimulus presentations. Further behavioral analyses revealed that the trial-unique object benefit was driven specifically by a reduction in false alarms rather than by enhanced hit rates. These findings indicate that the apparent advantage does not reflect expanded visual WM capacity per se, but rather the use of trial-unique stimuli that are less susceptible to PI.

Finally, hierarchical DPSD modeling of the empirical ROCs revealed a key dissociation between underlying recognition processes associated with the observed behavioral pattern. The recollection parameter, hypothesized to reflect context-bound WM representations, remained comparable across all stimulus type conditions, while familiarity reflecting context-free episodic LTM traces was selectively enhanced for trial-unique meaningful objects. Thus, the meaningful object advantage under trial-unique conditions appears predominantly to be attributed to enhanced familiarity signals provided by episodic LTM rather than expanded active WM storage capacity.

### Experiment 2

While the results of Experiment 1 suggest that the meaningful object advantage reflects reduced PI and enhanced contributions from LTM rather than expanded WM capacity, direct measurement of the number of neurally-active representations in WM provides a powerful complement to the behavioral evidence. Thus, Experiment 2 examined the CDA, a neural marker of active WM storage, in a lateralized change detection task involving trial-repeated colors and trial-unique meaningful objects across varying set sizes (1, 3, or 5 items). Here, we would like to explain our rationale for leaving out the trial-repeated condition from this EEG study. Our primary objective was to test whether prior reports of a behavioral advantage for meaningful objects was due to changes in online storage in WM, per se, rather than enhanced contributions from LTM. Although Experiment 1 provides one line of evidence for the latter explanation, it may still be possible to conceive of other explanations for the impact of repeating a small set of objects throughout the experiment. For instance, one might speculate that subjects in the trial-repeated condition were affected by semantic satiation (Smith & Klein, 1990), a phenomenon in which the semantic processing of a word diminishes with repeated presentations. If this had happened, then the absence of a behavioral benefit could be explained by the minimization of semantic content rather than by the imposition of high levels of PI. Given arguments of this kind, observing no difference in CDA amplitude between trial-repeated objects and colors would not provide clear evidence for our preferred explanation. By contrast, by including the trial-unique condition, we had the opportunity to replicate (once again) prior observations of the behavioral advantage for objects over colors, and then to directly observe whether online neural signatures of WM storage are affected by stimulus type. If a larger number of trial-unique objects can be stored in WM, then our design is well positioned to detect this effect. Thus, we focused on obtaining excellent statistical power for comparing WM storage of trial-unique objects and simple colors.

A critical prediction centers on the *interaction* between set size and stimulus type (Figure 5). If meaningful objects increase visual WM capacity, CDA amplitude should rise with set size and plateau at a higher level for meaningful objects than for simple features, producing a significant set size × stimulus type interaction. Alternatively, if the advantage reflects contributions from episodic familiarity under PI-free conditions, no change in the shape of the CDA by set size function is predicted. Importantly, by measuring CDA activity across multiple set sizes, we will be able to tell whether any observed differences in CDA activity reflect differences in WM storage (yielding an interaction between stimulus type and set size) or stimulus-driven effects that are independent of changes in the number of items stored in WM.

**Figure 5.**
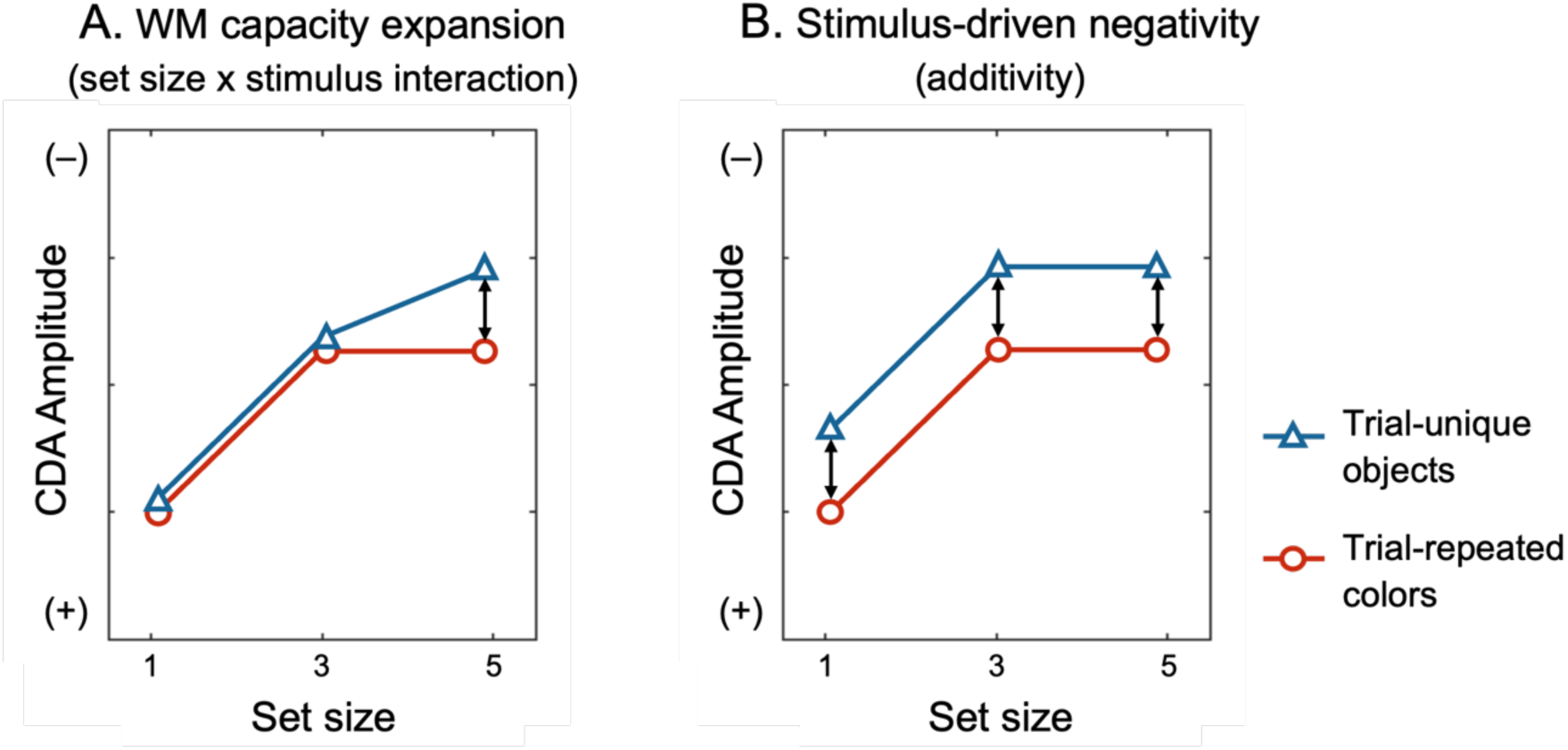
Hypothetical CDA amplitude patterns predicted by competing hypotheses. (A) If remembering meaningful objects truly expands visual WM capacity, the CDA amplitude should exhibit a significant set size × stimulus type interaction effect, with CDA amplitude increasing across a larger range of set sizes (set size 5). (B) By contrast, a stimulus-related confound hypothesis predicts only additive amplitude differences between stimulus types across all set sizes, without interaction.

## Method

### Participants

Twenty-five volunteers (14 female) participated in Experiment 2. All participants were right-handed, aged 19-32 years (*M* = 23.7, *SD* = 3.1), and reported normal or corrected-to-normal vision. Informed consent was obtained according to procedures approved by the University of Chicago Institutional Review Board. The final sample size of 18 participants and number of trials collected was determined based on prior EEG studies investigating CDA amplitude differences under various stimulus conditions, with an eye towards studies examining the CDA amplitude by set size function (Fukuda et al., 2015; Luria et al., 2016; Woodman & Vogel, 2008).

### Experimental Design

Participants completed a lateralized change detection task in two blocked conditions, trial-repeated colors and trial-unique objects, respectively (Figure 6A). Stimulus generation was identical to Experiment 1. Each trial began with a 500 ms central fixation (radius of 0.2°), followed by a 500 ms arrow cue presented above the fixation circle, indicating the side to remember (left/right, equally probable). A memory array was presented for 1,000 ms. In order to account for the lateralized nature of the CDA component, the memory array contained 1, 3, or 5 items presented in each visual hemifield. For both hemifields, the matching number of items were displayed at randomly chosen locations among five fixed placeholders (3.0° × 3.0° each), two were on an inner imaginary circle of 2.7° radius and the other three were on an outer imaginary circle of 7.0° radius, with 2.0° lateralized offset from the central midline of the display. After a 1000 ms retention interval displaying only placeholders, a test probe appeared at one of the remembered item locations on the cued side. Participants indicated whether the probe matched the original item by pressing one of two buttons, “z” or “/” key to indicate “no-change” or “change”, respectively. The probability of change and no-change was equal. Participants completed 200 trials per set size condition in each of the repeated-color and unique-object blocks with a randomized block order across participants.

**Figure 6.**
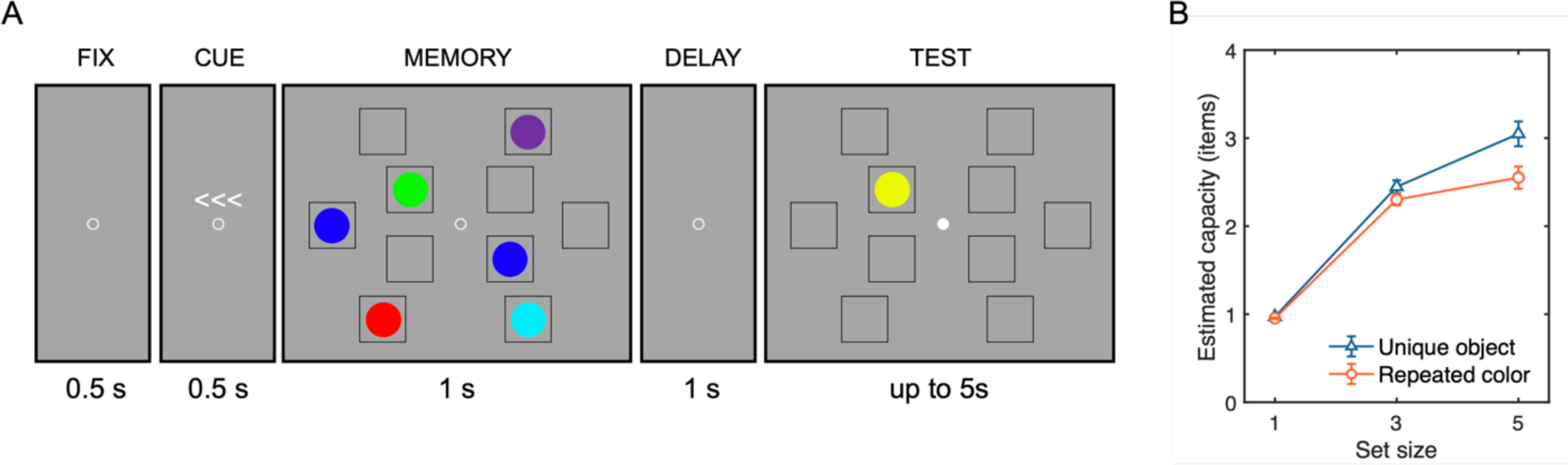
Procedure and resulting capacity estimates for Experiment 2. (A) Lateralized visual WM task sequence shown for the color stimulus condition. Each trial began with central fixation (500 ms), followed by an arrow cue indicating the task-relevant hemifield (left or right; 500 ms). The memory array was then presented laterally (1,000 ms), followed by a blank retention interval (1,500 ms). Participants subsequently indicated whether a test probe matched the remembered item at the corresponding location. The fixation point changed its color to white to indicate the onset of test array and allow participants to blink. The trial-unique meaningful object condition used the same procedure, differing only in stimulus content. (B) Capacity estimates of mean Cowan’s *K*, as a function of set size (1, 3, or 5 items) and stimulus type (unique object vs. repeated color). Error bars represent standard error of the mean.

### EEG Acquisition

EEG was recorded from 30 active Ag/AgCl electrodes (actiCHamp, Brain Products, Munich, Germany) mounted in an elastic cap positioned according to the international 10-20 system (Fp1, Fp2, F7, F3, F4, F8, Fz, FC5, FC6, FC1, FC2, C3, C4, Cz, CP5, CP6, CP1, CP2, P7, P8, P3, P4, Pz, PO7, PO8, PO3, PO4, O1, O2, Oz). A ground electrode was placed at position Fpz and two additional electrodes were affixed with stickers to the left and right mastoids. Data were referenced online to the right mastoid and re-referenced offline to the algebraic average of the left and right mastoids. Incoming data were filtered (low cutoff = 0.01 Hz, high cutoff = 80 Hz; slope from low to high cutoff = 12 dB/octave) and recorded with a 500 Hz sampling rate. Impedance values were kept below 10 kΩ.

Eye movements and blinks were monitored using electrooculogram (EOG) activity and eye tracking. We collected EOG data with five passive electrodes (two vertical VEOG electrodes placed above and below the right eye, two horizontal HEOG electrodes placed ∼1 cm from the outer canthi of each eye, and a ground electrode placed on the left cheek). Eye-tracking data was collected using a desk-mounted EyeLink 1000 Plus eye-tracking camera (SR Research, Ontario, Canada) sampling at 1,000 Hz.

### Artifact Rejection

EEG data were segmented into epochs (−200 to 2,000 ms from memory array onset). An automatic artifact rejection pipeline was applied to detect eye movements, blinks, and EEG artifacts. We implemented a comprehensive set of criteria using ERPLAB functions (Lopez-Calderon & Luck, 2014). Trials contaminated by artifacts were excluded for EEG analyses but not from behavioral analyses. We rejected trials containing eye movements and blinks using EOG channels and eye-tracking data.

For EOG channels, trials were flagged when the absolute voltage exceeded 50 μV or when step-like activity exceeded 30 μV within a 100 ms moving window (advanced in 10 ms steps). For eye-tracking rejection, we applied a similar sliding window to the x-gaze coordinates and y-gaze coordinates (window size = 100 ms, step size = 10 ms, threshold = 0.5°). When eye-tracking data were not available, we used EOGs to detect saccades and blinks. For EEG artifacts, we checked for drift (e.g., skin potentials) using the *pop_rejtrend* function, excluding trials in which a line fitted to the EEG data had a slope exceeding 75 μV with a minimal *R²* of 0.3. High-frequency noise and muscle artifacts were detected using the *pop_artmwppth* function, excluding trials with peak-to-peak activity greater than 75 μV within a 200-ms sliding window advanced in 100-ms steps. We also excluded trials with absolute voltage exceeding ±100 μV in any EEG channel using the *pop_artextval* function. Additionally, we detected step-like artifacts (which can occur with electrode movement) using the *pop_artstep* function, flagging trials with voltage changes exceeding 60 μV within a 150-ms sliding window advanced in 10-ms steps. For ocular artifacts, we applied separate criteria to EOG channels, using an absolute voltage threshold of ±50 μV and detecting saccade-like step functions exceeding 30 μV within a 100-ms sliding window advanced in 10-ms steps. Trials containing flatline signals were also excluded. Seven participants with rejection rate greater than 30% of trials were excluded from further analysis. Across the remaining participants, an average of 12.7% of trials were rejected, with no significant differences in rejection rates between stimulus conditions or set sizes. CDA was calculated as mean amplitude differences between contralateral and ipsilateral posterior electrodes (PO3/PO4, PO7/PO8, P3/P4, P7/P8; Figure 7). The average CDA amplitudes were taken from three different measurement windows, during stimulus presentation (i.e., encoding; 400–1,000 ms), delay period (1,400–2,000 ms), and a combined window of 400–2,000 ms following memory onset.

**Figure 7.**
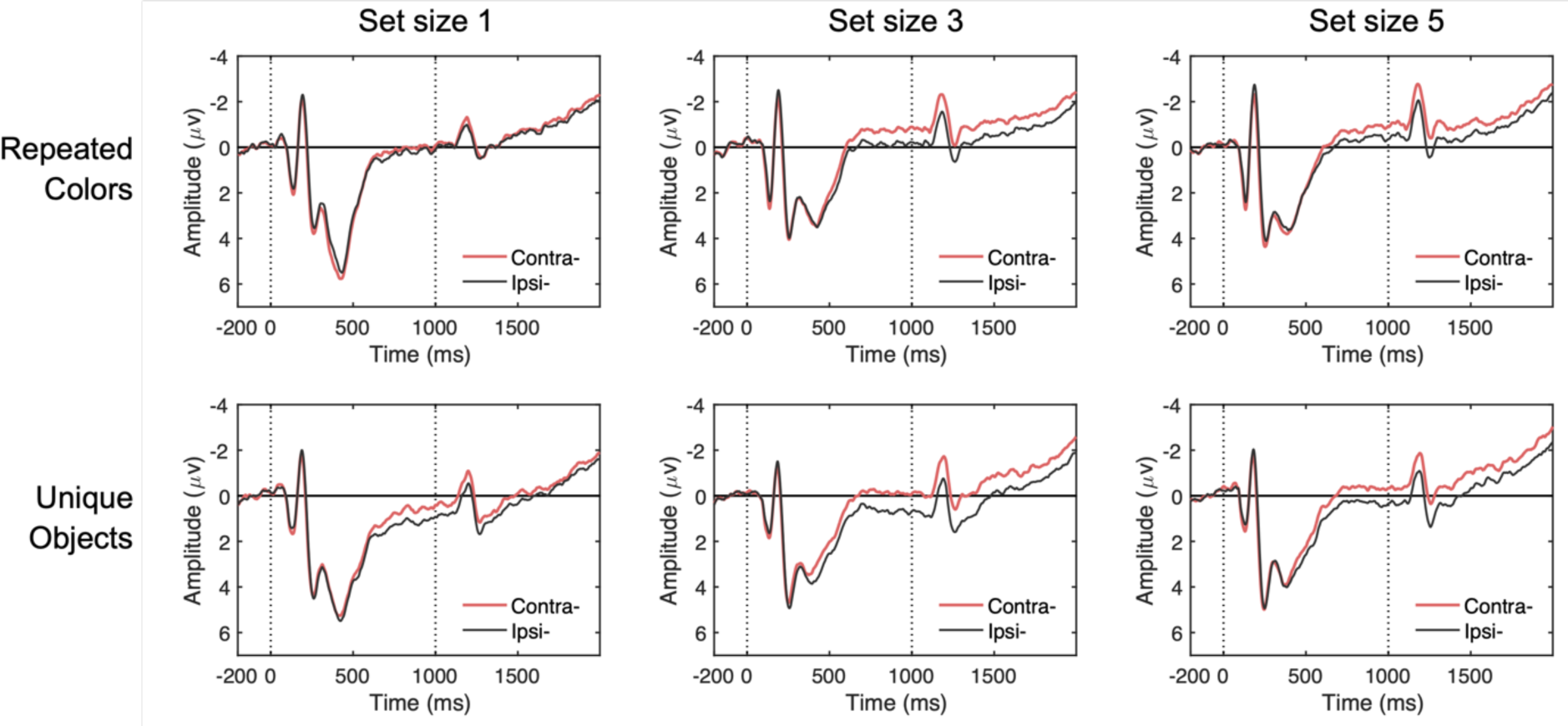
Grand-averaged contralateral and ipsilateral waveforms by stimulus type and set size. Waveforms are time-locked to the memory array onset (0 ms) and offset (1000 ms) and averaged across posterior electrode sites (P3/P4, P7/P8, PO3/PO4, PO7/PO8) in Experiment 2. Each panel compares contralateral (red) and ipsilateral (black) activity for trial-repeated colors (top row) versus trial-unique objects (bottom row) at set sizes 1, 3, and 5 (left to right). Vertical dotted lines mark the onset and offset of the memory array.

## Results

### Behavioral Performance

Behavioral results mirrored those of Experiment 1 (Figure 6B). A two-way repeated-measures ANOVA on Cowan’s *K* as a function of stimulus type (trial-repeated color vs. trial-unique object) and set size (1, 3, vs. 5 items) revealed significant main effects of stimulus type, *F*(1, 24) = 13.61, *p* = .001, *η^2^p* = .36, and set size, *F*(2, 48) = 46.28, *p* < .001, *η^2^p* = .91. The set size × stimulus type interaction effect was also significant, *F*(2, 48) = 12.76, *p* < .001, *η^2^p* = .35.

Notably, the interaction effect was primarily driven by a significant difference in the *K* estimate at set size 5, with trial-unique objects being higher capacity estimates (*M* = 3.05, *SD* = 0.70) compared to trial-repeated colors (*M* = 2.55, *SD* = 0.63), *t*(24) = 4.04, *pbonf* = .001, *d* = 0.82. The difference in the *K* estimates between stimulus type at set size 1 (for unique objects: *M* = 0.97, *SD* = 0.04; for repeated colors: *M* = 0.95, *SD* = 0.04) and set size 3 (for unique objects: *M* = 2.45, *SD* = 0.37; for repeated colors: *M* = 2.30, *SD* = 0.33) did not yield statistical significance, *ts*(24) < 2.45, *psbonf* > .066, *d* < 0.50. Thus, we obtained another clear replication of the behavioral advantage for trial-unique meaningful objects and simple colors. This puts us in a strong position to use CDA activity to determine whether this behavioral advantage is due to increasing online storage in WM rather than enhanced contributions from LTM that would not yield an interaction between CDA activity and set size.

### CDA Results

Figure 8 shows the grand-averaged CDA waveforms and mean amplitudes across different time windows. In the combined analysis window (Figure 8B, right), a two-way repeated-measures ANOVA with factors of stimulus type (trial-repeated color vs. trial-unique object) and set size (1, 3, 5 items) revealed significant main effects of stimulus type, *F*(1, 17) = 21.33, *p* < .001, *η^2^p* = .56, and set size, *F*(2, 34) = 19.42, *p* < .001, *η^2^p* = .53. Critically, the set size × stimulus type interaction was not significant, *F*(2, 34) = 0.28, *p* = .760, *η^2^p* = .02, indicating that although meaningful objects elicited larger CDA amplitudes overall, this effect was *additive* with set size, indicating that it reflected a *stimulus-driven* increase in contralateral negativity that was independent of the number of items stored in WM.

**Figure 8.**
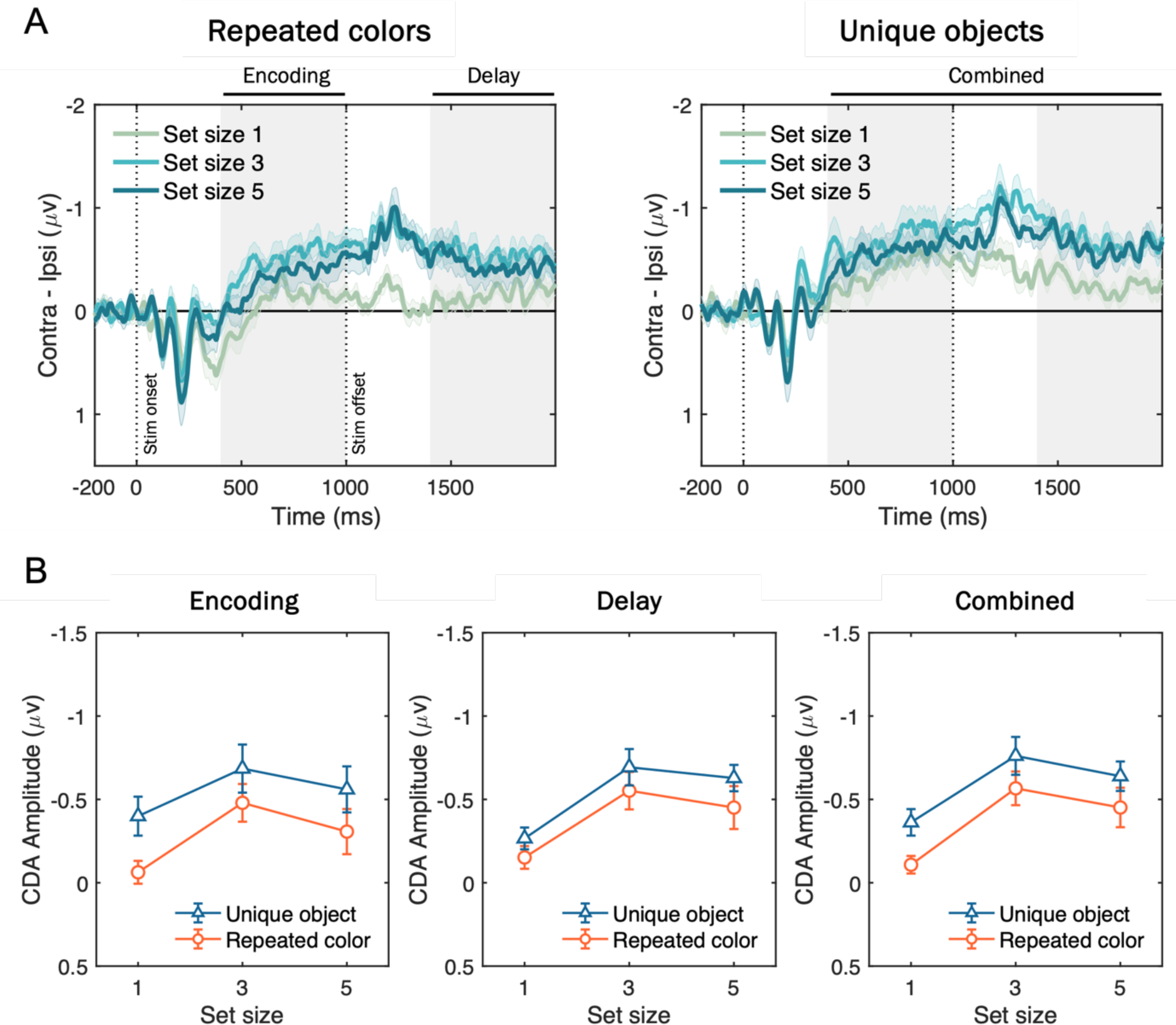
Grand-averaged CDA waveforms and mean amplitudes for trial-repeated colors and trial-unique objects. (A) CDA activity (contralateral minus ipsilateral activity at posterior electrode sites; P3/P4, P7/P8, PO3/PO4, PO7/PO8) time-locked to the memory onset and averaged across participants, plotted separately for repeated colors (left) and unique objects (right) at set sizes 1 (green), 3 (sky blue), and 5 (blue). Vertical dotted lines mark stimulus onset (0 ms) and offset (1000 ms), and gray shading indicates Encoding (400–1000 ms from stimulus onset), Delay (1400–2000 ms), and Combined (400–2000 ms) measurement windows. Shaded error bars represent standard error of the means (SEM). (B) Mean CDA amplitudes during the early (left), late (middle), and combined (right) time windows as a function of set size for unique objects (triangles on blue line) versus repeated colors (circles on red line). Error bars represent SEM.

These results patterns remained the same when analyzing the separate time windows during stimulus presentation (i.e., encoding; Figure 8B, left) or delay period (i.e., delay; Figure 8B, middle), both showing that the two-way interaction effects did not achieve statistical significance, *F*(2, 34) = 1.44, *p* = .250, *η^2^p* = .08 (encoding), and *F*(2, 34) = 0.01, *p* = .909, *η^2^p* = .01 (delay). Furthermore, including measurement window as a factor (encoding vs. delay) in a three-way ANOVA revealed no significant main effect of window, *F*(1, 17) = 0.17, *p* = .682, *η^2^p* = .01, nor a three-way interaction, *F*(2, 34) = 0.171, *p* = .843, *η^2^p* = .01. these findings demonstrate that while meaningful objects elicited larger CDA amplitudes than colored squares, the effect was purely additive and did not reflect an increase in active storage capacity.

## Discussion

Using an electrophysiological measure of neurally active storage in WM, Experiment 2 provided evidence against the claim that more meaningful than simple objects can be stored in visual WM. Although behavioral data showed superior performance for trial-unique meaningful objects over repeated colors, the shape of the CDA by set size function revealed that the same number of items were stored in WM. If subjects were able to store more meaningful objects than simple colors, the difference in CDA amplitudes should have been larger for set size 5 where the clearest behavioral benefits were observed for meaningful objects over repeated colors. Instead, CDA differences between meaningful and simple objects were equivalent across all set sizes, suggesting a stimulus-driven contralateral negativity that is independent of the number of items stored in WM (Drew et al., 2011; Woodman & Vogel, 2008). Thus, online neural measures of WM storage disconfirm the hypothesis that WM capacity is expanded for meaningful objects.

## General Discussion

We present behavioral and neural evidence that calls into question recent claims that WM capacity is expanded for meaningful compared to simple objects. In Experiment 1, we manipulated meaningfulness and PI independently, and found that the performance advantage for meaningful stimuli was driven by trial-unique stimulus presentations that minimized PI rather than by meaningfulness per se. When PI was equated by using repeated meaningful stimuli, capacity estimates for meaningful and simple stimuli converged. Hierarchical Bayesian DPSD modeling further showed that the trial-unique advantage specifically reflected enhanced familiarity signals from episodic LTM, while recollection-based WM processes remained stable across stimulus types. Thus, improved performance under trial-unique conditions can be explained by increased LTM contributions when PI is minimized, rather than by increased WM storage capacity.

Experiment 2 provided converging electrophysiological evidence using the CDA, a neural marker of active WM maintenance. Several prior reports have described higher CDA amplitudes for meaningful objects than for simple features and have taken this as support for greater storage (Brady et al., 2016; Chung et al., 2024; Thibeault et al., 2024; Torres et al., 2024). However, those studies typically employed only a single set size, which cannot distinguish changes in the number of items stored from stimulus-driven differences in CDA amplitude that are independent of WM load. If meaningfulness truly increases the number of concurrently stored items, the CDA should exhibit an interaction between stimulus type and set size, with steeper increases and a plateau at higher set sizes for meaningful objects. Instead, the effect of meaningfulness on CDA was additive with the number of items to be stored, indicating a stimulus-related difference that is separable from load-dependent storage (Drew et al., 2011; Woodman & Vogel, 2008). This pattern aligns with demonstrations that some stimulus classes elicit larger contralateral negativities without altering the shape of the CDA-set size function, the signature of changes in WM storage.

Note that the CDA analysis in Experiment 2 focused on trial-unique meaningful objects and repeated colors and did not include repeated meaningful objects. This selection targeted the condition where a meaningfulness advantage was proposed, testing whether the CDA-set size function shows an interaction with stimulus type. Including a repeated meaningful object condition would add little inferential power because Experiment 1 showed that equating PI yielded comparable behavioral capacity and stable recollection parameters for repeated meaningful objects and repeated colors. Under those conditions, the anticipated CDA outcome for repeated meaningful objects would be an overlapping set size function with colors, with any additive offset that is not diagnostic of storage, consistent with prior demonstrations of additive stimulus effects (Drew et al., 2011; Woodman & Vogel, 2008).

From a broader theoretical perspective, these findings align closely with embedded-process models of WM (Cowan, 2001; Oberauer, 2002). In these accounts, a capacity-limited focus of attention operates over activated LTM representations, maintaining a small set of individuated items in an immediately accessible state via attentional pointers that support binding to spatiotemporal context (Awh & Vogel, 2025; Oberauer & Lin, 2017). Within this framework, meaningfulness can enhance encoding quality and decision evidence through access to LTM, yet the number of items concurrently maintained by attentional pointers remains limited. Therefore, performance benefits observed for meaningful stimuli under trial-unique conditions reflect reliance on episodic familiarity signals from LTM, in the absence of any change in the number of actively maintained items.

WM capacity has often been described as item based rather than feature based. Classic findings showed similar capacity for single feature and multi feature objects, which supports object-based limits (Luck & Vogel, 1997). Later work found modest costs for multi-feature objects but reliable “object-based benefits”, with features stored more effectively when bound within a single item (Hardman & Cowan, 2015; Ngiam et al., 2024; Olson & Jiang, 2002). The present results are consistent with that view. Capacity estimates converged when interference was equated, and the CDA-set size functions did not interact with stimulus type. These properties indicate stable item limits despite differences in stimulus richness. Relatedly, associative chunking in LTM can yield WM performance that resembles storing more items without increasing the number of concurrently maintained units. Learned word pairs can perform like single items, and consistently paired colors can resemble single colors in WM tasks (Chen & Cowan, 2009; Brady et al., 2009). These benefits depend on retrievable LTM representations, rather than an expansion of WM storage (Ngiam et al., 2019; Huang & Awh, 2018). This perspective dovetails with the present DPSD results, where recollection remained stable while familiarity increased under trial-unique conditions.

Our study highlights PI as a critical gating mechanism that regulates interactions between episodic LTM and active WM. Under low-PI conditions (e.g., trial-unique stimuli), episodic LTM provides reliable, context-free familiarity signals that enhance performance. Conversely, high-PI conditions (e.g., repeated stimuli) make episodic familiarity an unreliable cue to “oldness”, forcing greater reliance on context-specific recollection supported by active WM where representations are bound to spatiotemporal pointers (Awh & Vogel, 2025). This gating clarifies why traditional WM paradigms with repeated stimuli yield stable capacity estimates around three or four items, whereas trial-unique paradigms can yield much larger apparent capacities (Endress & Potter, 2014). The inflation arises because familiarity boosts recognition without increasing the number of online representations. Thus, differences between trial-unique and repeated designs are not simply methodological details but determine whether behavioral performance reflects the capacity-limited WM or familiarity-based LTM contributions.

An additional theoretical consideration is semantic satiation, in which repeated exposure to meaningful stimuli temporarily reduces semantic distinctiveness and can impair recognition (Smith & Klein, 1990; Tian & Huber, 2010). If satiation were the primary cause of reduced performance with repeated meaningful objects, deficits should be selective to meaningful stimuli and align with electrophysiological indices of semantic access rather than storage. Two observations argue against this account. First, when interference was equated by repeating a fixed stimulus set, performance converged for meaningful objects and for abstract colors lacking lexical-semantic associations, implicating PI rather than semantic loss. Second, in Experiment 2 with trial-unique meaningful objects, where satiation should be minimal, the CDA difference between meaningful objects and colors was additive across set sizes and showed no set-size interaction. Thus, while satiation may exacerbate familiarity reductions for meaningful stimuli under heavy repetition, it does not explain the full pattern: matched capacity under repetition, stable recollection parameters, and additive CDA offsets without load interactions. Future work can test a satiation contribution by pairing repetition counts that induce satiation with N400 measures of semantic access and CDA set-size functions to determine whether repetition alters access without altering storage.

Our findings connect to evidence supporting hierarchical interactions between memory systems, comprising perceptual processes and contextual binding mechanisms (Christophel et al., 2017; Lee et al., 2023; Yatziv & Kessler, 2018). Perceptual memory provides large-capacity but interference-prone representations automatically encoded into activated LTM, whereas active WM engages selective binding to maintain individuated items in context. Recent neuroimaging and neurophysiological findings are consistent with this dual-code perspective, with sensory-based representations supported by posterior cortices and abstract, context-bound codes supported by frontoparietal networks (Chunharas et al., 2024; Kwak & Curtis, 2022; Xu & Chun, 2006). Intracranial recordings similarly dissociate medial temporal neurons that encode detailed sensory and semantic features from frontal neurons associated with abstract, slot-like WM capacities (Kamiński et al., 2017). In this view, meaningfulness enhances the strength or distinctiveness of sensory representations and boosts familiarity, but does not change the number of items that can be indexed by the attentional pointer system.

Several limitations suggest promising directions for future research. For example, while we controlled PI through stimulus repetition, we did not systematically manipulate graded levels of PI or examine its spatial specificity (Endress, 2022; Donenfeld et al., 2024; Lin & Luck, 2012; Makovski, 2016). In principle, one possibility is that familiarity-driven benefits may scale monotonically with PI reduction across trials. Investigating the temporal dynamics of PI build-up and release (Keppel & Underwood, 1962; Watkins & Watkins, 1975) could further clarify how global familiarity-based LTM processes can shape memory performance in WM procedures.

Moreover, our findings should be considered in light of growing research on memorability, a stimulus-intrinsic property that makes certain images consistently better remembered (Bainbridge, 2019; Isola et al., 2014). Recent work shows that intrinsic memorability effects also emerge during visual WM tasks (Tam et al., 2025; Ye et al., 2024). Future studies exploring interactions between PI and memorability may elucidate how stimulus properties and task contexts jointly influence memory performance, potentially integrating stimulus-centered and process-based accounts.

In conclusion, our behavioral and neural data demonstrate that previously reported WM capacity advantages for meaningful objects are better explained by reduced PI and elevated episodic familiarity than by expansions of active storage capacity. These findings reinforce theoretical frameworks in which a fixed-capacity attentional pointer mechanism limits the number of concurrently maintained items, while LTM contributes variable decision evidence depending on interference (Bartsch et al., 2024; Oberauer & Bartsch, 2023). By decoupling meaningfulness from PI and by measuring full CDA set-size functions, future studies can avoid misattributing familiarity-based gains to storage and can provide sharper tests of WM capacity limits.

## Acknowledgments

This work was supported by the National Institute of Mental Health (ROIMH087214) and Office of Naval Research (N00014-12-1-0972) awarded to E.A.

## Footnotes

1 Even if a study employed a continuous ring of colors in a recall procedure, it is well known that observers perceive a small set of discrete categories within this color space (e.g., Bae et al., 2015)

2 The one exception to the use of a single set size is the study by Brady and colleagues (2016). However, there has been continued debate about the precise empirical pattern, because subsequent replication attempts have not been successful. Quirk and colleagues (2020) ran a replication study using the same stimuli and task, but did not observe CDA evidence for increased storage of meaningful objects. Moreover, Brady and colleagues (2016) reported that performance with meaningful objects benefitted more from increased exposure durations than did simple colors, but this effect was not replicated either. Given uncertainty regarding the robustness of these findings, we focus our discussion on the other published studies that examine CDA activity with meaningful and simple objects.

